# Sex-Specific Neurovascular and Cognitive Deficits in Offspring of Preeclampsia

**DOI:** 10.1101/2025.08.21.671659

**Authors:** Felipe Troncoso, Hermes Sandoval, Esthefanny Escudero-Guevara, Francisco Nualart, Eder Ramírez, Pedro Sandaña, Joakim Ek, Owen Herrock, Daymara Merceron, Álvaro O. Ardiles, Manu Vatish, Jesenia Acurio, Carlos Escudero

## Abstract

**Background:** Offspring of preeclampsia may develop long-term neurovascular and cognitive impairments, but the underlying mechanisms remain unclear. We investigate sex-specific alterations in brain vascular function and cognition in a murine model of preeclampsia induced by the nitric oxide synthase inhibitor L-NAME.

**Methods:** Pregnant mice received L-NAME (gestational day 7 to 19). Offspring were assessed at postnatal day 5 (P5) and in adulthood (4–5 months). Brain vascular development, angiogenesis, perfusion, cold-induced vasoconstriction, and blood-brain barrier (BBB) integrity were assessed in vivo and ex vivo. Offspring’s serum was applied to brain endothelial cell cultures to evaluate endothelial activation and barrier function. Adult cognitive performance was evaluated through behavioral tests and hippocampal long-term potentiation (LTP) recordings.

**Results:** L-NAME offspring (P5) exhibited reduced brain vascular density, decreased tip cell formation, and impaired cold-induced vasoconstriction, most prominently in males. BBB integrity was compromised, with increased permeability and reduced expression of tight junction proteins (claudin-5, ZO-1), again more pronounced in males. Serum from L-NAME offspring induced endothelial activation and barrier dysfunction in vitro. These effects were accompanied by increased brain and systemic levels of hypoxia markers and proinflammatory cytokines (IL-6, TNF-α). Adult L-NAME offspring showed cognitive deficits in recognition and spatial memory tasks, with female offspring displaying more pronounced impairments. These behavioral findings paralleled a reduced magnitude and impaired stabilization of hippocampal LTP in females.

**Conclusions:** Prenatal exposure to a preeclampsia-like environment induces persistent, sex-specific neurovascular and cognitive deficits in offspring. These findings underscore the need for long-term neurological follow-up in children born to preeclamptic pregnancies.

## Introduction

Preeclampsia is a severe hypertensive disorder of pregnancy characterized by placental dysfunction, new-onset hypertension after 20 weeks of gestation, and signs of maternal organ damage, including proteinuria and neurological involvement^1-4^. It poses significant risks for both mother and fetus. Infants born to preeclamptic pregnancies have an increased risk of perinatal morbidity and mortality^5,6^, with male infants appearing particularly vulnerable^7,8^. Preclinical studies have similarly shown that male offspring of preeclampsia-like animal models are more susceptible to cardiovascular alterations than females^9,10^.

The fetal brain is susceptible to adverse intrauterine conditions. Children born to mothers with preeclampsia are at heightened risk for perinatal stroke^11-13^, neonatal encephalopathy^14,15^, and long-term cognitive impairments associated with altered brain anatomy, connectivity, and perfusion^16-19^. In a recent postmortem study, we reported increased cerebrovascular vulnerability in fetuses and neonates exposed to maternal hypertension, including a higher incidence of subarachnoid hemorrhage^20^. These findings emphasize the significance of cerebrovascular integrity in early life for maintaining long-term brain health^21,22^.

Furthermore, sex differences have been reported in cerebral blood flow (CBF) in neonates, in whom males may have higher CBF in the parietal, frontal, and temporal lobes^23^. Understanding how pregnancy complications such as preeclampsia affect these physiological differences is crucial. However, data on CBF and vascular function in offspring of preeclamptic pregnancies remain scarce. Only one small case-control study reported lower CBF in children born to women with preeclampsia, but without examining sex differences^17^.

Sex-specific differences have emerged as a relevant aspect of preeclampsia-related outcomes in cerebrovascular function. While some studies report a higher incidence of brain injury in male offspring^24,25^, both sexes appear to be affected. We previously found that neonates (postnatal day 5, P5) from a preeclampsia-like model characterized by reduced uterine perfusion pressure (RUPP) displayed greater signs of brain edema than their counterparts in the control group. Notably, brain perfusion deficits were more pronounced in female RUPP-offspring^26^. These findings highlight the sexually dimorphic nature of cerebrovascular vulnerability following prenatal hypertensive exposure.

Animal models have played a crucial role in elucidating how preeclampsia affects brain development in the offspring. The L-NAME-induced preeclampsia-like model, based on nitric oxide synthase inhibition, mimics several features of human preeclampsia, including maternal hypertension, proteinuria, and fetal growth restriction^27^. This model has revealed impaired neurogenesis, reduced brain angiogenesis, and cognitive deficits in exposed offspring^27-30^. However, the functional status of cerebral blood vessels— specifically their vasoreactivity and blood-brain barrier (BBB) properties—remains poorly characterized in this context.

Therefore, in this study, we hypothesized that offspring exposed to maternal preeclampsia-like conditions would exhibit sex-specific alterations in the BBB integrity and brain endothelial activation, leading to divergent cognitive outcomes in adulthood.

## Methods

The data supporting this study’s findings are available from the corresponding author. Further methodological details are provided in the Supplementary Information and associated references^26,27,31-41^.

### Statistical analysis

Quantitative variables are presented as median ± standard deviation, whereas qualitative variables are presented as percentages within their respective groups. Considering data distribution, we used parametric or non-parametric tests, as appropriate, using the Shapiro-Wilk normality test. Then, we compared the groups by Student’s t-test or Mann-Whitney test. In the analysis of brain homogenates and *in vitro* studies, where experiments were performed head-to-head, a paired t-test was used. p<0.05 was considered a statistically significant difference. Data and statistical analyses were performed using a Microsoft Excel database and GraphPad Prism 6 (GraphPad Software, CA, USA).

## Results

### Offspring from preeclampsia-like syndrome showed reduced brain angiogenesis

Our group has previously characterized the preeclampsia-like syndrome induced by L-NAME administration^27^. At postnatal day 5 (P5), offspring from L-NAME-treated dams were smaller than controls, regardless of sex (Table S1).

We have previously shown reduced brain angiogenesis in offspring from L-NAME dams at P5^27^. Here, we investigated whether this effect exhibited sex-specific patterns. Compared to controls, offspring from L-NAME dams exhibited significantly lower vascular density (Figure S1A), shorter vessel length (Figure S1B), reduced expression of GLUT1, a marker of brain endothelial cells (Figure S1C), and fewer tip endothelial cells (Figure S1D) in the brain cortex. These reductions were evident in both male and female offspring from L-NAME dams (Figure 1A–1E). However, only female offspring from L-NAME dams displayed significantly lower vascular density than female controls (Figure 1D).

**Figure 1.**
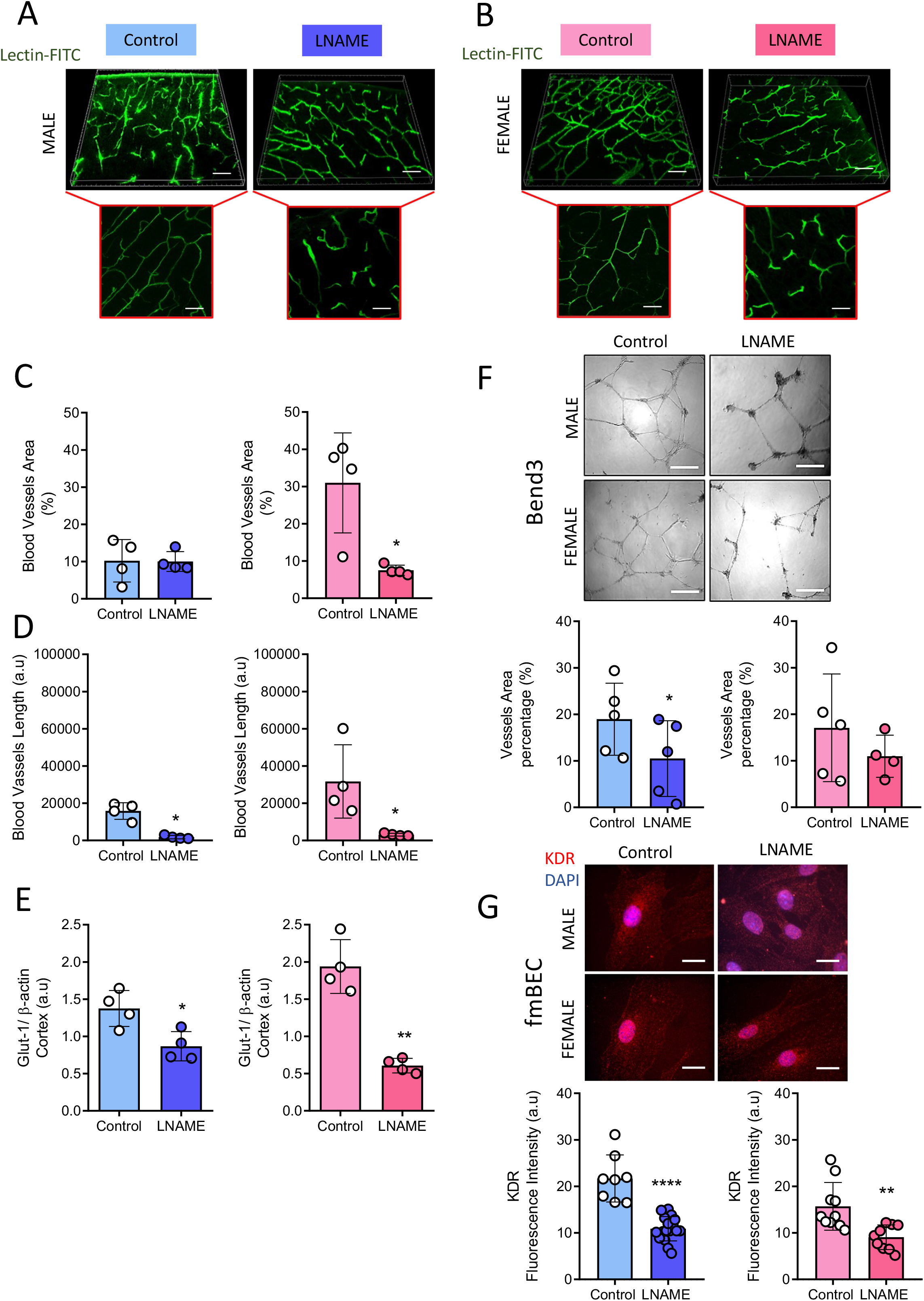
Reduced brain angiogenesis in male and female offspring from preeclampsia-like syndrome. A-B) 3D reconstructions of cortical vasculature labeled with fluorescent lectin in male (A) and female (B) pups from control (Control) and L-NAME-treated dams (LNAME). Bottom panels: 2D magnifications. C-D) Quantification of cortical vascular density (C) and vessel length (D) in male (blue bars) and female (pink bars) P5 pups. E) Densitometric analysis of GLUT-1 (brain endothelial marker) normalized to β-actin in cortical homogenates from male and female P5 pups. F) Representative images of Bend3 cells treated with serum from male and female P5 pups. Graphs indicate quantification of *in vitro* angiogenic capacity as vessel area in comparative groups. G) Representative immunofluorescence images of KDR (VEGFR2) and its quantification in fetal brain endothelial cells (fmBEC) exposed to serum from male and female P5 pups. Each dot represents a randomly selected individual subject. Data are presented as median ± standard deviation. *p < 0.05, **p < 0.005, ****p < 0.0001 vs. respective control group.

Given previous reports of dysregulated circulating factors impairing endothelial angiogenic capacity in offspring exposed to preeclampsia^27,42^ we confirmed this effect in a female-derived brain endothelial cell line (Bend3) treated with serum (50 μg/ml, 3 h) from P5 offspring of L-NAME-treated dams (Figure S1E–S1F). Further analysis revealed that serum from male—but not female—offspring from L-NAME-treated dams significantly reduced angiogenic capacity in Bend3, as determined by reduction in vessel area and the number of vessel junctions (Figures 1F; Figure S1G).

Given that Bend3 cells are derived from adult female mouse brain endothelium, we next isolated primary brain endothelial cells from male and female fetuses of normal pregnancies and exposed them to sex-matched serum from offspring of control or L-NAME dams. Compared to controls, serum from L-NAME offspring significantly reduced KDR protein levels (the critical proangiogenic receptor) in these primary fetal brain endothelial cultures (fmBEC, Figure S1H), with similar reductions observed in both male- and female-derived cells (Figure 1G), indicating a sex-independent impairment of proangiogenic signaling.

### Offspring from L-NAME dams showed reduced cold-induced brain microvascular vasoconstriction

To evaluate whether the reduced angiogenesis observed in offspring from L-NAME dams affected brain perfusion, we assessed cerebral microvascular blood flow under basal conditions and following cold-induced vasoconstriction (Figure S2A-B). Baseline brain perfusion was similar between L-NAME and control offspring (Figure S2C). However, following exposure to a cold stimulus, offspring from L-NAME dams failed to show the expected vasoconstrictive response (Figure S2C–S2D). This impairment was significant only in male offspring, who failed to mount an appropriate vasoreactive response, whereas female offspring showed no significant difference from female controls (Figure S2E–S2F).

### Offspring from L-NAME showed an impaired blood-brain barrier

One critical characteristic of the brain blood vessels is their function as a barrier, i.e., forming the BBB. To assess BBB integrity, we analyzed cortical endothelial markers. Compared to controls, offspring from L-NAME dams exhibited increased expression of CD31 (also known as PECAM-1), a marker of endothelial activation (Figure S3A), with this elevation significant only in male offspring (Figure 2A-2B).

**Figure 2.**
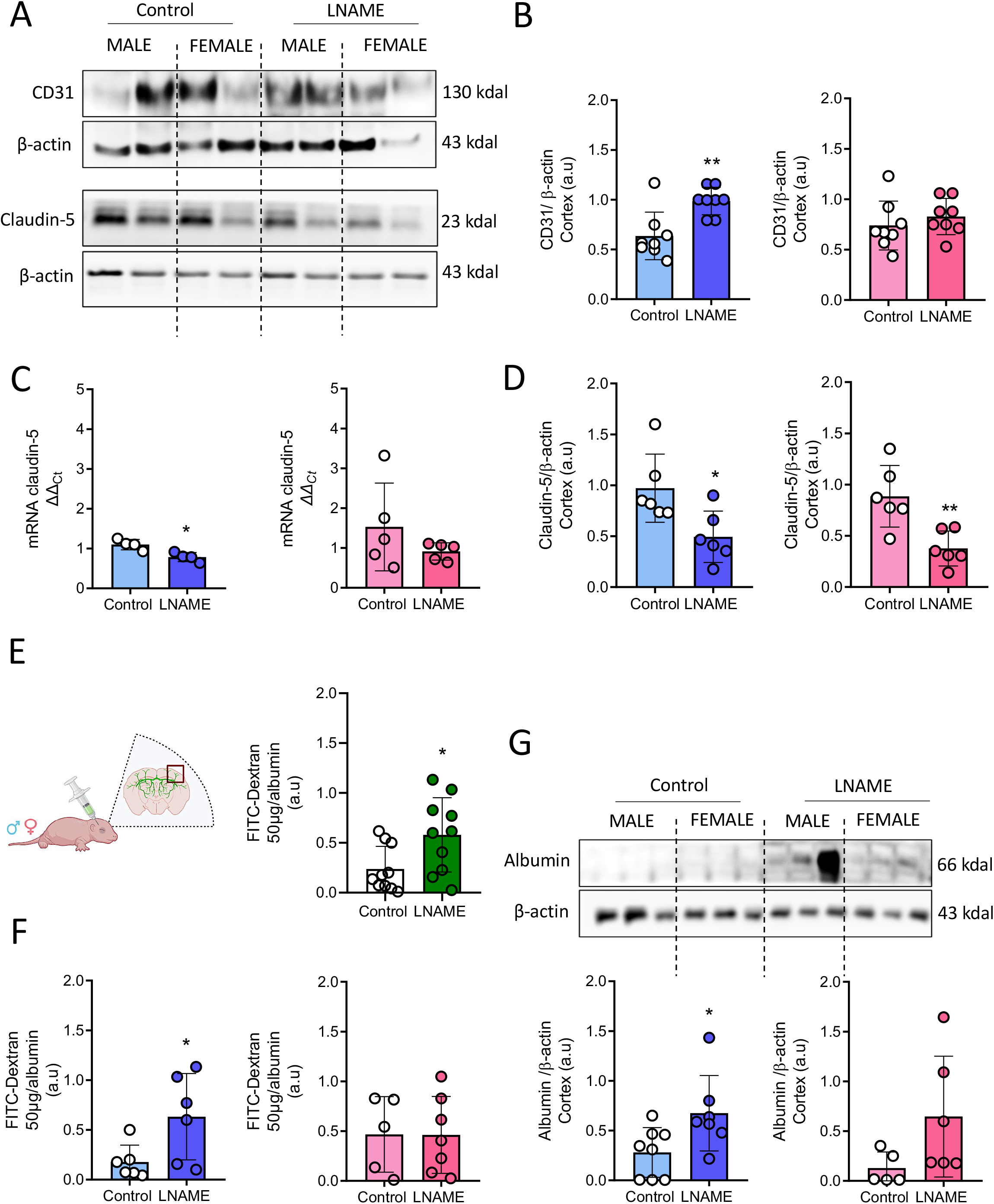
Disruption of the blood-brain barrier in male and female offspring of preeclampsia. A) Representative blots of the endothelial marker CD31and CLDN5 in brain cortex homogenates from P5 pups of control (Control) and L-NAME (LNAME) dams. B) Densitometric analysis of CD31 normalized to β-actin in cortical homogenates from male (blue bars) and female (pink bars) P5 pups. C-D) mRNA (C) and protein (D) levels of the tight junction protein Claudin 5 in cortical homogenates from male and female P5 pups. E) Schematic of the experimental approach with intraocular injection of FITC-Dextran-70 kDa. Cartoon was created with BioRender. Graphs indicate quantification of FITC-Dextran-70 kDa extravasation in cortical homogenates from all subjects, with separate analyses in F) male (blue bars) and female (pink bars) pups. G) Representative blot of albumin extravasation in brain cortex homogenates from control and L-NAME pups (male and female); β-actin served as loading control. Graphs indicate densitometric analysis of albumin/β-actin ratio as indicated in representative blots. Each dot represents a randomly selected individual subject. Data are presented as median ± standard deviation. *p < 0.05, **p < 0.005 vs. respective control group.

In contrast, both mRNA and protein levels of CLDN5, a tight junction protein essential for BBB function^43^, were significantly reduced in offspring from L-NAME dams compared to controls (Figure S3B-C). This reduction was observed in both sexes (Figure 2C-2D).

We then assessed BBB permeability *in vivo* (Figure 2E) using FITC-dextran extravasation. L-NAME offspring displayed significantly higher cortical FITC-dextran accumulation than controls (Figure 2E), indicating increased BBB permeability. This effect was significantly pronounced in male offspring from L-NAME dams (Figure 2F).

To confirm barrier disruption, we evaluated albumin levels in cortical homogenates via Western blot. Albumin, a plasma protein excluded by an intact BBB, was significantly elevated in L-NAME offspring compared to controls (Figure 2G, Figure S3D), with this increase driven predominantly by male offspring (Figure 2G).

### Brain hypoxic and inflammatory response in offspring from L-NAME dams

To assess the brain’s response to vascular impairment, we measured levels of hypoxia-inducible factor 1-alpha (HIF-1α) and the proinflammatory cytokines, interleukin-6 (IL-6) and tumor necrosis factor-alpha (TNF-α) in the cortex of P5 offspring from L-NAME dams (Figure 3A). Compared to controls, offspring from L-NAME dams showed increased protein levels of HIF-1α (Figure S4A), as well as elevated IL-6 at both the mRNA and protein levels (Figure S4B-S4C). However, TNF-α protein levels, but not mRNA levels, were reduced in this group (Figure S4D-S4E).

**Figure 3.**
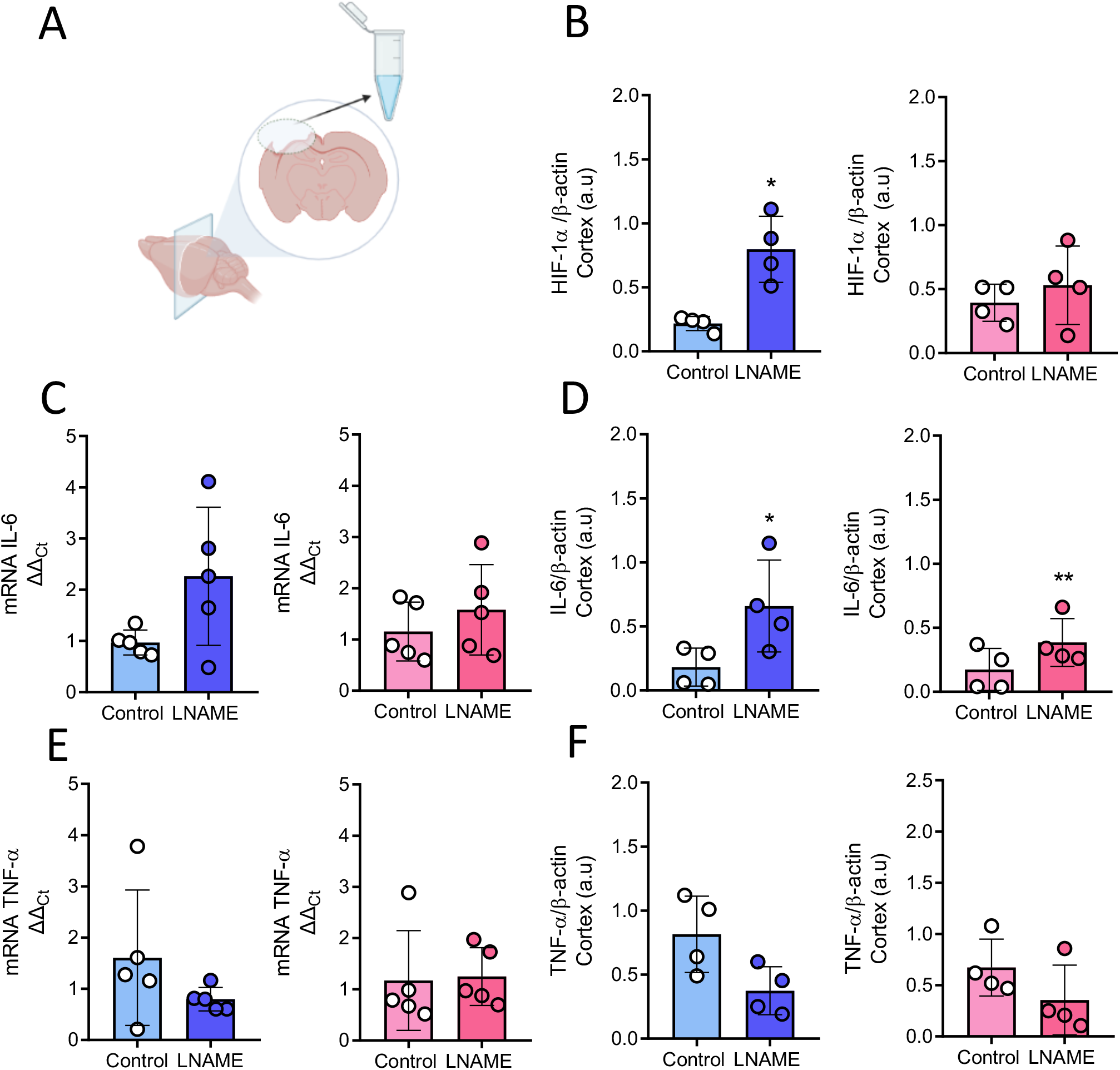
Proinflammatory profile in the brains of male and female offspring of preeclamptic mothers. A) Schematic of brain cortex homogenate preparation. Cartoon was created with BioRender. B) Densitometric analysis of HIF-1α/β-actin ratio as a hypoxia marker in male (blue bars) and female (pink bars) P5 pups from control (Control) and L-NAME dams (LNAME). C-D) mRNA (C) and protein (D) levels of IL-6 in the brain cortex as in A. E-F) mRNA (E) and protein (F) levels of TNF-α as in A. Each dot represents a randomly selected individual subject. Data are presented as median ± standard deviation. *p < 0.05, **p < 0.005 vs. respective control group.

Sex-stratified analysis revealed that elevations in HIF-1α and IL-6 protein levels remained statistically significant in male offspring of L-NAME treated dams compared to their counterparts in the control group (Figure 3B–3D). However, in female L-NAME offspring, only IL-6 protein levels were significantly increased (Figure 3D). We found no changes in the synthesis of TNF-α at both the mRNA and protein levels in male and female offspring between the L-NAME and control groups (Figure 3E-3F).

We next assessed whether the proinflammatory profile observed in the brain cortex was reflected systemically (Figure 4A). Circulating levels of IL-6 (Figure S5A) and IL-8 (Figure S5B) were elevated in offspring from L-NAME-treated dams compared to controls. TNF-α levels, despite high variability, did not differ significantly between groups (Figure S5C).

**Figure 4.**
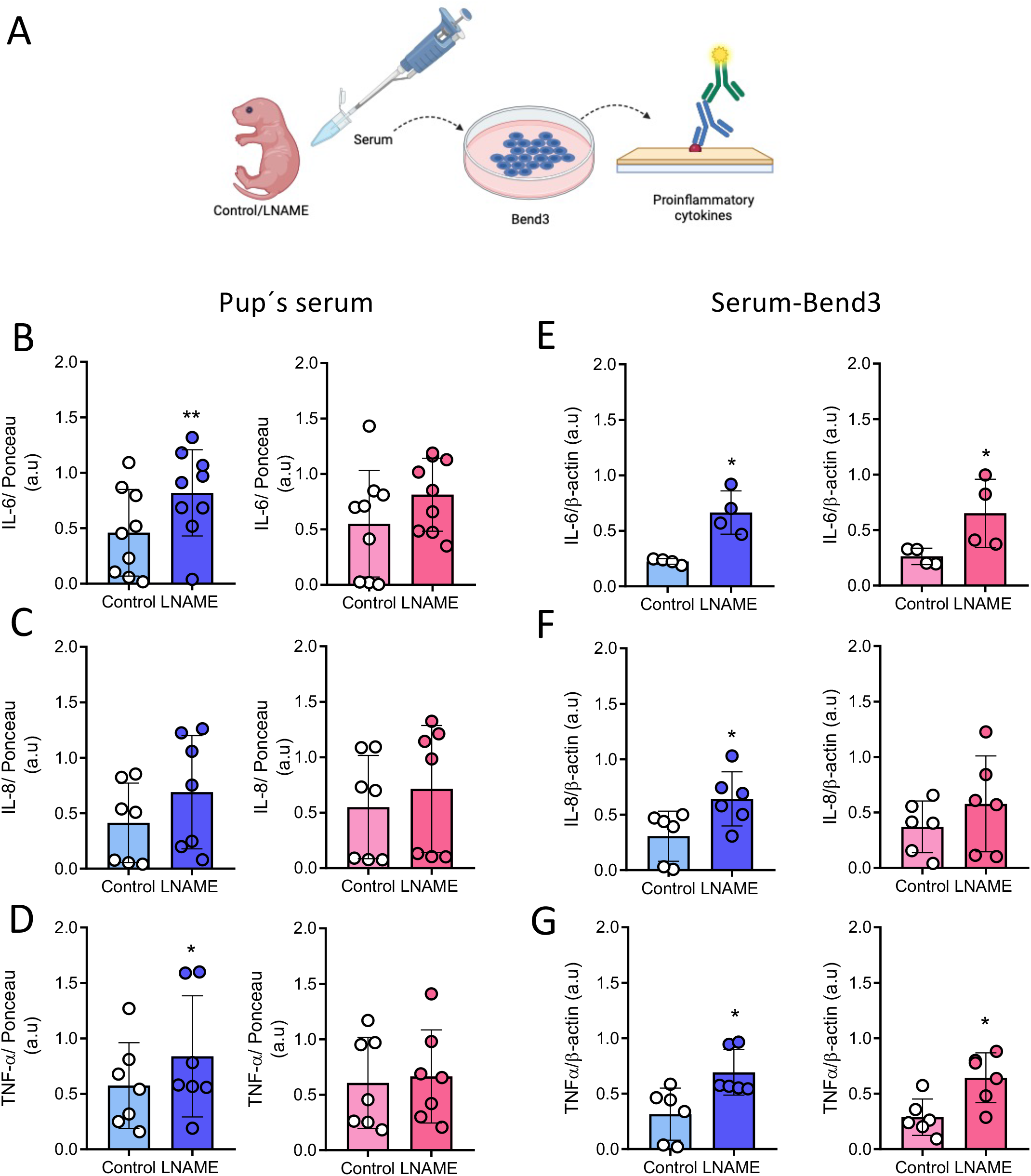
Proinflammatory serum profile in male and female offspring from preeclampsia and brain endothelial cell activation. A) Schematic showing serum collection from P5 pups of control (Control) and L-NAME dams (LNAME). Cartoon was created with BioRender. Serum was analyzed for circulating levels of B) IL-6, C) IL-8, and D) TNF-α using Western blot in male (blue bars) and female (pink bars) pups. Serum from male (blue bars) and female (pink bars) pups was also used to stimulate mouse brain endothelial cells (Bend3) to evaluate protein levels of E) IL-6, F) IL-8, G) TNF-α by Western blot. Each dot represents a randomly selected individual subject. Data are presented as median ± standard deviation. *p < 0.05, **p < 0.005 vs. respective control group.

Stratification by sex revealed that circulating levels of IL-6 and TNF-α were significantly elevated in male offspring from L-NAME-treated dams compared to male controls (Figure 4B, 4D). In contrast, the previously observed differences in IL-8 levels did not remain significant following sex-specific analysis (Figure 4C). These findings suggest a partial overlap between systemic and cerebral inflammatory responses, with stronger systemic changes in male offspring.

### Serum from offspring of L-NAME dams triggered a proinflammatory response in brain endothelial cells

To assess whether circulating factors from L-NAME-exposed offspring could activate brain endothelial cells, we treated Bend3 cells with serum from P5 pups (Figure 4A). Compared to control serum, serum from L-NAME-treated offspring induced significantly higher protein levels of IL-6 (Figure S5D), IL-8 (Figure S5E), TNF-α (Figure S5F), and the endothelial activation marker CD31 (Figure S5G).

Sex-stratified analysis showed that IL-6 (Figure 4E) and TNF-α (Figure 4G) synthesis remained significantly elevated in Bend3 cells treated with serum from both male and female offspring of L-NAME dams. Conversely, IL-8 synthesis was significantly increased only in cells treated with serum from male L-NAME offspring (Figure 4F). At the same time, the previously observed increase in CD31 synthesis was not significant after stratifying by sex (data not shown). These results suggest that serum from L-NAME offspring—particularly males—can directly trigger a proinflammatory response in brain endothelial cells.

We next examined whether serum from L-NAME offspring affected BBB function *in vitro* (Figure 5A). Compared to controls, serum from L-NAME offspring significantly reduced transendothelial electrical resistance (TEER; Figure S6A), increased permeability to FITC-Dextran (Figure S6B), and decreased protein levels of the tight junction proteins, CLDN5 (Figure S5C) and zonula occludens-1 (ZO-1; Figure S6D).

**Figure 5.**
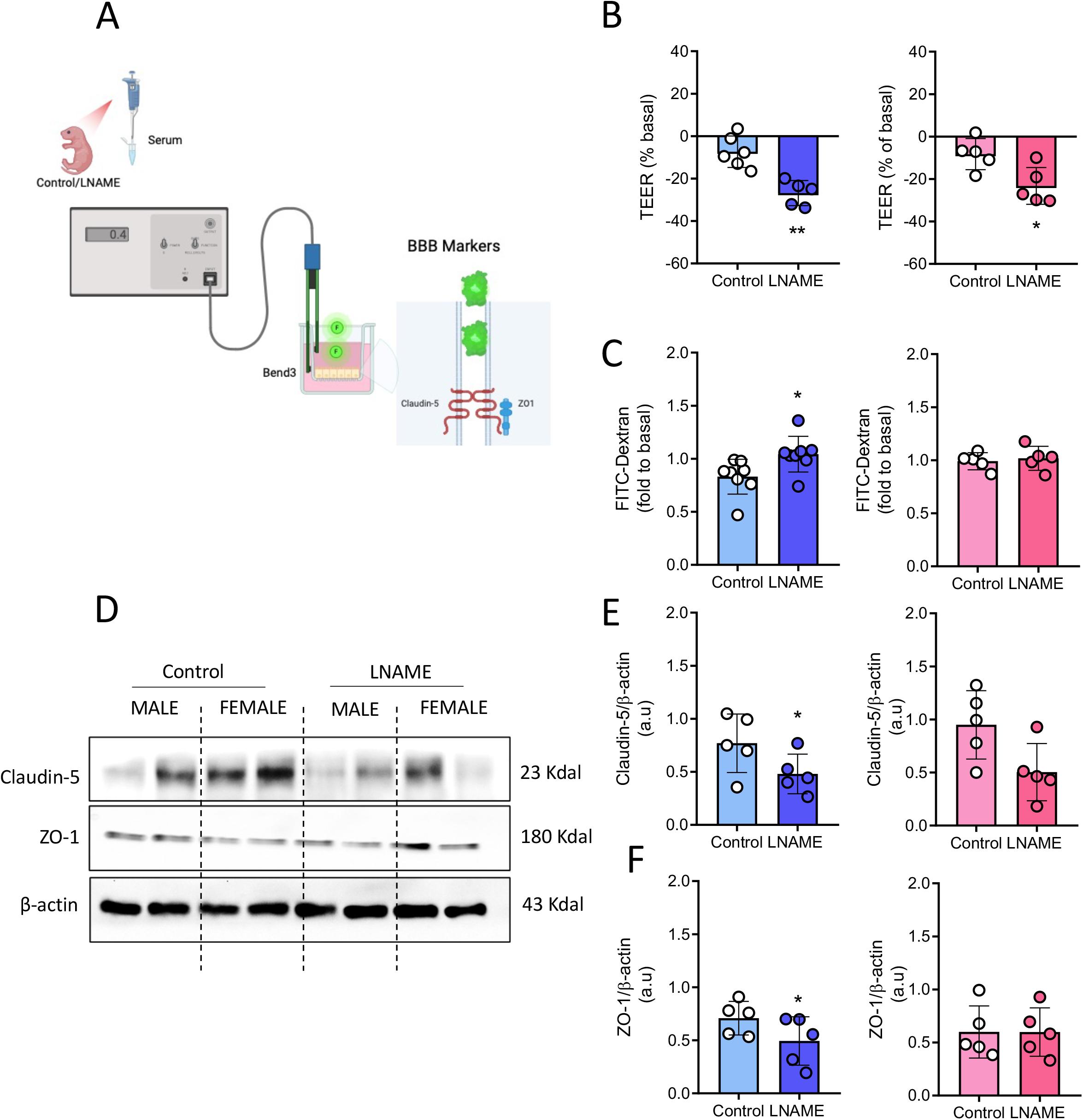
Serum from male offspring of preeclamptic mothers and barrier properties of Bend3. A) Schematic showing serum from P5 male (blue bars) and female (pink bars) offspring of control (Control) and L-NAME dams (LNAME) used to stimulate Bend3 cells and assess barrier function. Cartoon was created with BioRender. B) Percentage change in transendothelial electrical resistance (TEER) and C) permeability to FITC-Dextran (70 kDa) in Bend3 cells treated with serum from male and female pups of control and L-NAME dams. D) Representative blots of tight junction proteins (Claudin-5 and ZO-1) and loading control β-actin in Bend3 cells treated as in A. E) Densitometric analysis of Claudin-5/β-actin and F) ZO-1/β-actin ratios in Bend3 cells. Each dot represents a randomly selected individual subject. Data are presented as median ± standard deviation. *p < 0.05, **p < 0.005 vs. respective control group.

When analyzed by sex, significant impairment of all four BBB parameters was observed only in cells treated with serum from male L-NAME offspring (Figure 5B-5F). However, serum from female L-NAME offspring still induced a significant drop in the electrical resistance (Figure 5B), suggesting partial barrier dysfunction. These results indicate that sex-specific circulating factors contribute to BBB disruption in offspring exposed to a preeclampsia-like environment.

To validate these findings from Bend3 cells in primary cultures of brain endothelial cells, we treated fetal mouse brain endothelial cells (fmBEC) with serum from L-NAME and control offspring (Figure 6). Serum from offspring of L-NAME dams significantly increased CD31 protein expression, indicating endothelial activation (Figure S7A). This effect was statistically significant only in cells treated with serum from male offspring (Figure 6A-6B).

**Figure 6.**
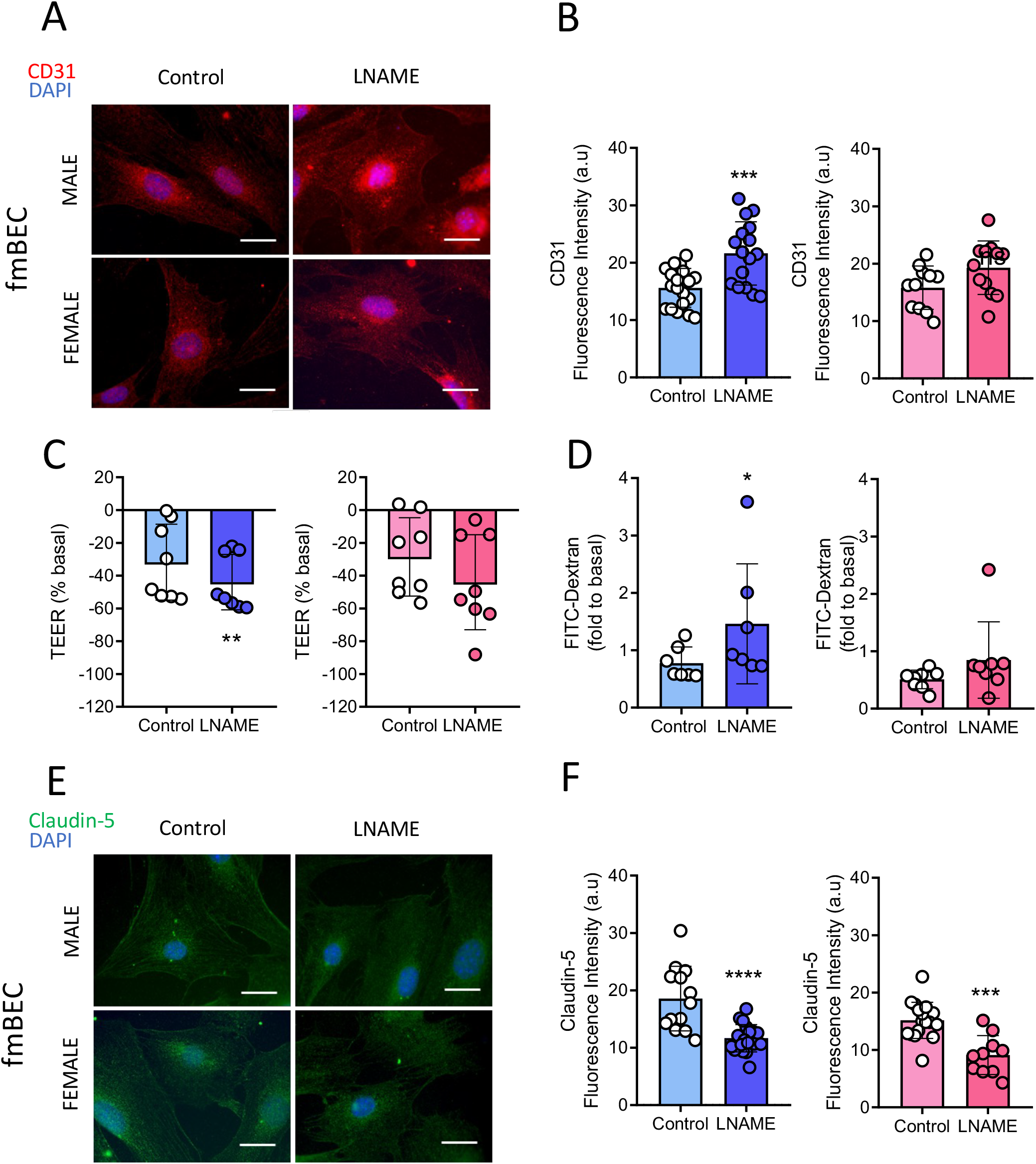
Serum from offspring of preeclampsia and barrier properties of primary fetal mouse brain endothelial cells. A) Representative images of immunodetection of the endothelial activation marker CD31 in primary fetal mouse brain endothelial cells (fmBEC) exposed to serum from P5 male (blue bars) and female (pink bars) offspring of control (Control) and L-NAME dams (LNAME). B) Quantification of CD31 immunodetection, C) percentage change in transendothelial electrical resistance (TEER), and D) permeability to FITC-Dextran (70 kDa) in fmBEC treated as in A. E) Representative images of claudin-5 immunodetection and F) quantification of claudin-5 immunofluorescence in fmBEC treated as in A. Each dot represents a randomly selected individual subject. Data are presented as median ± standard deviation. *p < 0.05, **p < 0.005, ***p < 0.01, ****p < 0.0001 vs. respective control group.

Consistent with findings in the adult brain endothelial cell line Bend3 (Figure 5), serum from L-NAME offspring impaired barrier function in fmBEC, as evidenced by decreased TEER, increased FITC-Dextran permeability, and reduced CLDN5 protein expression (Figure S7B–S7D). Notably, barrier disruption remained significant in fmBEC treated with serum from male offspring, as shown by reduced TEER (Figure 6C) and increased permeability (Figure 6D). In contrast, CLDN5 expression was similarly decreased following exposure to serum from both male and female offspring (Figure 6E–6F), suggesting that some aspects of barrier disruption are sex-independent.

### Adult consequences of impaired brain vascular function in offspring from L-NAME dams

To evaluate long-term consequences of brain vascular alterations, cognitive performance was assessed in adult offspring from L-NAME-treated dams using four behavioral tests: Open Field Test (OFT), Novel Object Recognition Test (NORT), Place Recognition Test (PRT), and Morris Water Maze Test (MWMT) (Figure S8A). No significant differences were observed in total distance traveled (Figure S8B) or locomotor velocity (Figure S8C) in the OFT, indicating comparable baseline motor activity across groups.

In contrast, in the NORT (Figure S8D) and PRT (Figure S8E), control offspring showed a significant preference for the novel object or location (N), respectively, over the familiar one (F), reflecting intact recognition and spatial memory. This preference was absent in L-NAME offspring, suggesting impaired hippocampal-dependent memory. Similarly, in the MWMT (Figure S8F), L-NAME offspring exhibited reduced time in the target (NW) quadrant where the hidden platform was located. Conversely, they showed an increased time in the opposite (SE) quadrant compared to control offspring, further indicating spatial learning deficits.

Sex-stratified analysis showed that offspring from L-NAME-treated dams had fewer crossings in the OFT compared to controls, with similar reductions in both males and females (Figure 7A). In the MWMT, offspring of L-NAME dams exhibited increased escape latency during the acquisition phase in both sexes (Figure 7B). This impairment was evident throughout the four-day analysis in females, whereas in males it became significant only during the third and fourth days (Figure 7B). These findings indicate that prenatal exposure to L-NAME impairs both locomotor activity and spatial learning in offspring, with cognitive deficits emerging earlier and more persistently in females than in males.

**Figure 7.**
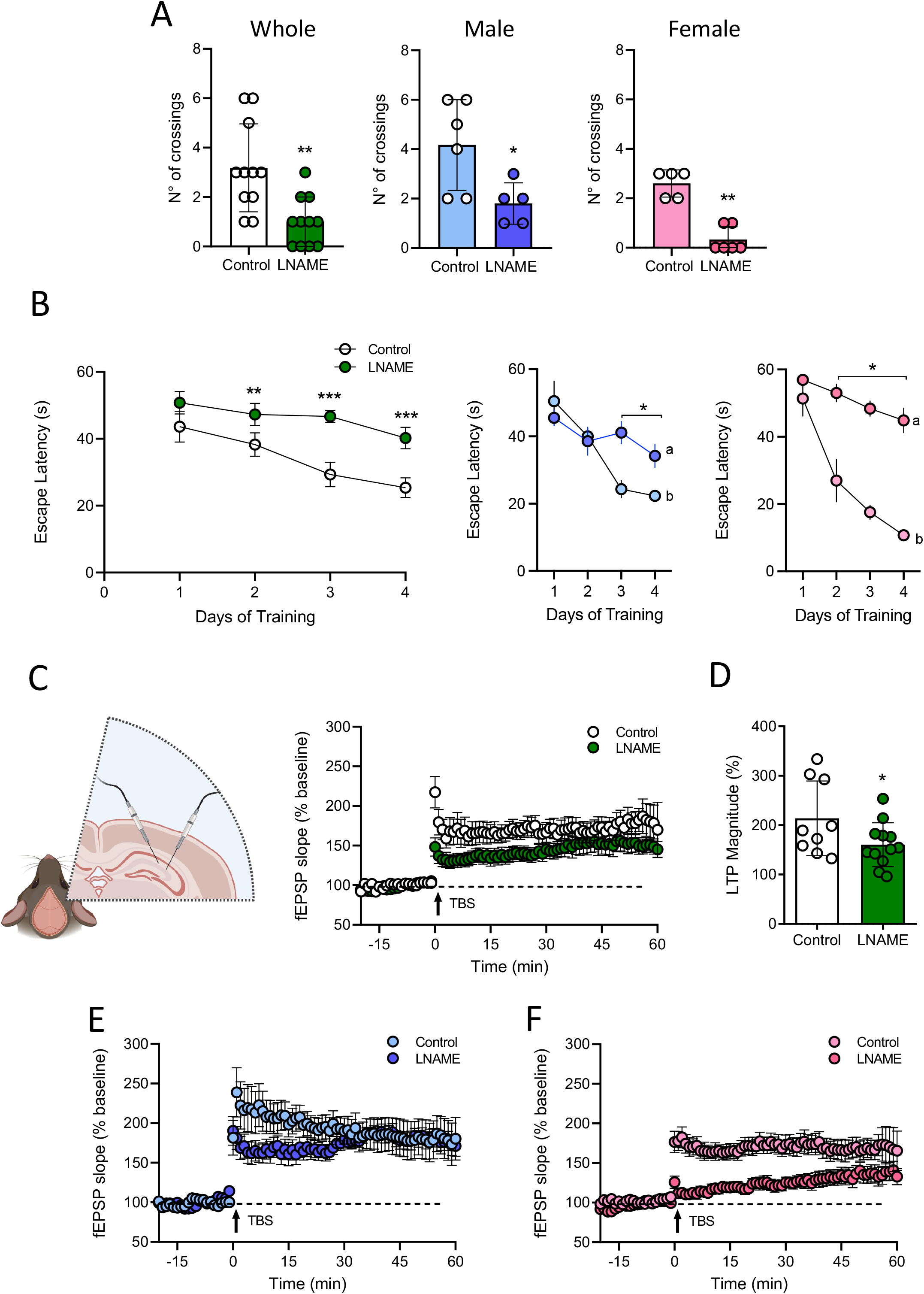
Spatial learning and synaptic plasticity in adult offspring of preeclampsia. Adult offspring (4-5 months) of control (Control) and L-NAME-treated dams (LNAME) underwent behavioral testing and hippocampal electrophysiological analysis. A) Number of crossings in the target quadrant during the probe trial of the Morris water maze, analyzed in the entire offspring group (Whole) or separated by sex. Male offspring (blue bars) and female offspring (pink bars) from control and L-NAME groups. B) Escape latency during acquisition sessions across all offspring. As well as being separated by sex as indicated in A. C) Cartoon of hippocampal electrophysiological analysis and representative field excitatory postsynaptic potential (fEPSP) traces recorded from hippocampal slices of the entire group.Cartoon was created with BioRender. TBS, theta burst stimulation. D) Long-term potentiation (LTP) magnitude, expressed as percent change from baseline, in hippocampal slices from the same offspring groups as in A. E-F) Representative fEPSP traces recorded in hippocampal slices from male (E) and female (F) offspring of control and L-NAME dams. Each dot represents a randomly selected individual subject. Data are presented as median ± standard deviation. *p < 0.05, **p < 0.005, ***p < 0.01 vs. respective control group.

### Long-Term Potentiation (LTP) in offspring from L-NAME dams

To assess synaptic plasticity in adult offspring from L-NAME dams, *ex vivo* hippocampal long-term potentiation (LTP) recordings were performed in brain slices from male and female mice. Group-level analysis showed a reduced magnitude of LTP in offspring from L-NAME-treated dams compared to controls (Figure 7C-7D). When stratified by sex, female offspring from L-NAME-treated dams displayed a diminished and gradually increasing postsynaptic response following induction, indicative of impaired LTP stabilization (Figure 7F). In contrast, male L-NAME offspring showed LTP responses comparable to controls (Figure 7E).

To explore whether synaptic function was linked to behavioral outcomes, LTP magnitude was correlated with performance in the PRT and MWMT (Figure S8H–S8I). In control offspring, greater LTP was positively associated with better cognitive performance. However, this relationship was absent in L-NAME offspring, suggesting a decoupling between hippocampal plasticity and memory function following prenatal exposure to a preeclampsia-like environment.

## Discussion

Offspring from preeclampsia-like syndrome generated by maternal L-NAME administration exhibited both structural (e.g., reduced vascular density, tip cells, GLUT1) and functional (e.g., cold-induced vasoconstriction, BBB integrity) deficits of the brain blood vessels. These changes affect both sexes but are more pronounced in males. Elevated cortical levels of HIF-1α and IL-6 suggest a hypoxic, proinflammatory brain environment, consistent with prior reports of neuroinflammation in this context^44,45^. Prenatal L-NAME exposure impairs locomotor activity and spatial learning in both sexes, but with earlier effects in females, consistent with their selective deficit in hippocampal LTP stabilization. Collectively, this study demonstrates that intrauterine exposure to a preeclampsia-like environment induces long-lasting, sex-specific alterations in brain vascular development, BBB integrity, and cognitive function in the offspring.

While sex-specific effects on the developing brain following preeclampsia are increasingly recognized, data on sex differences in cerebrovascular outcomes remain limited. Preeclampsia is associated with a twofold increased risk of perinatal stroke^11,46^, with male neonates disproportionately affected. Male infants are generally more vulnerable to adverse perinatal outcomes, including severe asphyxia^47^, cerebral palsy ^48^, or greater long-term IQ impairment than female infants with a similar degree of hypoxia-induced encephalopathy^49^. This sex-specific vulnerability is also reported in neonatal rodent models of hypoxia-ischemia injury ischemia injury^49-52^, as well as the fact that bacteremia exacerbates only males exposed to perinatal hypoxia-ischemia brain injury^53^. Animal studies further support a heightened susceptibility in males to long-term cardiovascular^9,10^ and cerebrovascular complications^24,25^. These findings are consistent with our current results, in which male and female offspring from preeclampsia exhibited cerebrovascular dysfunction, including angiogenesis and BBB disruption, with more consistent and pronounced effects observed in males.

The mechanisms driving impaired brain vascular development in offspring exposed to preeclampsia remain incompletely understood. Our findings show that serum from L-NAME offspring can disrupt angiogenesis and BBB integrity in brain endothelial cells, suggesting that circulating factors contribute to long-lasting cerebrovascular alterations. Previous work has shown that female HUVECs derived from preeclamptic pregnancies exhibit broader gene dysregulation and greater TNF-α–induced barrier disruption than male counterparts^54^, indicating also a potential role for sex-specific endothelial responses. These data support a model in which inflammatory mediators, potentially including proinflammatory cytokines, induce or exacerbate brain endothelial dysfunction in a sex-dependent manner.

In this regard, our data show that serum from preeclampsia-exposed offspring induces a proinflammatory phenotype in brain endothelial cells, with evidence of sex differences in circulating mediators. IL-6 and IL-8 were elevated in L-NAME offspring, but only IL-6 remained significantly increased in males, suggesting a potential role in sex-specific endothelial dysfunction. Although animal models consistently report elevated brain IL-6 after preeclampsia^44,45^, human data remain limited and inconsistent^55,56^. Notably, IL-6 can directly activate brain endothelium via JAK/STAT3 signaling in adult mice^57^. These findings highlight IL-6 as a candidate mediator of long-term neurovascular alterations in offspring from preeclampsia, warranting further investigation.

### Endothelial dysfunction and BBB impairment

We demonstrate consistent evidence of BBB disruption in offspring from preeclampsia-like pregnancies, with increased brain accumulation of fluorescent tracers and albumin, particularly in males. These effects were observed in both the RUPP^26^ and L-NAME models, and were replicated *in vitro* using primary brain endothelial cells. Reduced expression of tight junction proteins CLDN5 and ZO-1 further supports a mechanistic link to barrier breakdown. This BBB leakiness, together with impaired angiogenesis and blunted vascular reactivity, may contribute to the hypoxic and proinflammatory brain environment observed in exposed offspring. In turn, hypoxia may generate a feedforward mechanism to exacerbate cerebrovascular dysfunction by further impairing vasodilatory signaling and angiogenesis^58,59^, and worsening BBB dysfunction^60^, as reported in models of stroke and neurodegeneration^61^.

### Long-term effects on cognition and synaptic plasticity

Adult offspring from preeclamptic pregnancies displayed hippocampal-dependent cognitive impairments, with no evidence of motor dysfunction. Although both sexes were affected in learning and memory tasks, female offspring showed more pronounced deficits, suggesting a heightened vulnerability. These results extend previous studies reporting reduced locomotor learning performance in female offspring from preeclamptic pregnancies^62^. Moreover, *ex vivo* hippocampal recordings revealed impaired synaptic plasticity in L-NAME offspring. While LTP correlated with behavioral performance in control animals, this relationship was absent in offspring exposed to preeclampsia, indicating a decoupling between synaptic plasticity and cognition. Collectively, these findings support the hypothesis that prenatal exposure to preeclampsia disrupts neurodevelopmental programming of the hippocampus, with long-term consequences for memory function. Intriguingly, the more substantial cognitive impairments observed in females may reflect a differential early-life adaptation to cerebrovascular dysfunction, warranting further investigation.

We acknowledge several limitations. Although we identified early cerebrovascular alterations in offspring of preeclampsia, we did not perform mechanistic experiments to define the circulating factors responsible. Vascular changes were more evident in males, whereas cognitive deficits predominated in females, a pattern that highlights the complexity and dynamic nature of neurovascular regulation rather than a binary contradiction. Our cross-sectional design and the analysis of different brain regions (cortex for vascular function vs. hippocampus for cognition) further limit direct interpretation. Future longitudinal and region-specific studies are required to clarify sex-dependent mechanisms linking vascular dysfunction and cognitive outcomes in this context.

In summary, prenatal exposure to L-NAME-induced preeclampsia causes early cerebrovascular dysfunction (including BBB leakiness, impaired angiogenesis, and reduced vascular reactivity) that extends into adulthood as cognitive impairment and hippocampal synaptic dysfunction. While male offspring exhibit more pronounced acute brain vascular impairment, females exhibit greater cognitive vulnerability later in life. These findings emphasize the role of early vascular programming in brain development and support the need to identify molecular targets for early diagnosis and intervention in offspring affected by preeclampsia.

## Perspective

The underlying mechanism of brain structure^16,17^ and neuronal connectivity^18,19^ present in children or pups of preeclamptic pregnancies, or their potential sex-dependent differences, is largely unknown. However, in hypoxic pups, preclinical findings indicate that male-associated disadvantages over female offspring include less functional and structural development of the lungs and cardiorespiratory circulation^63^; high circulating levels of testosterone associated with impaired stress response of the neonatal brain^50^; greater pro-apoptotic signaling pathways and neuronal cell death^50^; or less expression of the mitochondrial biogenesis-associated transcription factor, Nrf2/GABPα, leading to decreased electron transport chain proteins, and low antioxidants levels^51,52^. However, a critical actor, less studied in the literature, is the requirement of adequate cerebral blood flow (CBF) to maintain brain function.

In this context, our findings show that preeclampsia induces long-lasting, sex-specific effects on the neurovascular system. Male offspring demonstrate more pronounced endothelial activation and BBB disruption, whereas female offspring exhibit greater vulnerability to cognitive impairment. These divergent outcomes may reflect intrinsic sex differences in immune programming, vascular maturation, or hormonal responsiveness during early development.

Our findings suggest that circulating inflammatory and anti-angiogenic mediators may serve as biomarkers of neurodevelopmental risk in offspring exposed to preeclampsia. The persistent endothelial activation induced by offspring serum suggests that the postnatal environment remains altered, prolonged after the initial in utero insult. Investigating the molecular pathways underlying these vascular and cognitive impairments may yield therapeutic targets to mitigate the long-term effects of maternal preeclampsia on offspring health. We encourage future follow-up studies in children exposed to preeclampsia using cognitive analysis and structural and functional MRI studies that allow us to identify vulnerabilities that require an early intervention.

## Abbreviations

(ACSF): Artificial cerebrospinal fluid
(BBB): Blood-brain barrier
(CBF): Cerebral blood flow
(CLDN5): Claudin 5
(Dx): Days of gestation
(fmBEC): Fetal mouse brain endothelial cells
(fEPSP): Field excitatory postsynaptic potentials
(HIF-1α): Hypoxia inducible factor 1 alfa
(hCMEC/D3): Human cerebral microvasculature endothelium
(IL-6): Interleukin-6
(IL-8): Interleukin-8
(KDR): Kinase insert-domain-containing receptor or Vascular endothelial growth factor receptor 2
(LTP): Long-term potentiation
(MCA): Middle cerebral artery
(MWM): Morris Water Maze Test
(Bend3): Murine brain endothelial cell line
(L-NAME): NG-Nitroarginine methyl ester hydrochloride
(NORT): Novel Object Recognition Test
(OFT): Open Field Test
(PRT): Place Recognition Test
(PlGF): Placental growth factor
(P5): Postnatal day 5
(fmBEC): Primary Fetal Mice Brain Endothelial Cell
(RUPP): Reduction of uterine perfusion
(ROIs): Regions of interest
(sFLT1, a decoy receptor of VEGF): Soluble fms-like tyrosine kinase
(TOI): Time of interest
(TNF-α): Tumor necrosis factor-alpha
(VEGF): Vascular endothelial growth factor
(ZO1): Zonula occludent 1

## Acknowledgment

The authors thank the Vascular Physiology Laboratory and GRIVAS Health researchers for their valuable input. This study was funded by Fondecyt 1200250 and GI2301146. 2023-2026 (Chile).

## Author contributions

CE conceptualized the manuscript. FT performed most of the experiments and analyses. JA, HS, and EEG performed some animal and *in vitro* experiments. FN and ER performed an analysis of brain angiogenesis by immunohistochemistry. PS was a clinician, pathologist, and consultant, and supported histological and immunohistochemical analysis. JE and OH perform additional brain vascular analysis by histochemistry. DM performs cognitive tests and electrophysiology experiments. AA supervises cognitive test analysis and performs electrophysiology experiments. MV advises on the clinical relevance of the findings and supports the experimental design. JA monitored the progress of experimental work. All co-authors approved the final version of this manuscript.

## Use of AI Tools

Portions of this manuscript’s writing were supported by OpenAI’s ChatGPT (GPT-4, July 2025 version) to improve clarity, language, and coherence. The authors critically reviewed, edited, and approved all AI-assisted content. No AI tools were used for data analysis, interpretation of results, or generation of scientific conclusions.

